# β-arrestin mediates communication between plasma membrane and intracellular GPCRs to regulate signaling

**DOI:** 10.1101/2020.04.08.031542

**Authors:** Maxwell S. DeNies, Alan Smrcka, Santiago Schnell, Allen P. Liu

## Abstract

It has become increasingly apparent that G protein-coupled receptor (GPCR) localization is a master regulator of cell signaling. However, the molecular mechanisms involved in this process are not well understood. To date, observations of intracellular GPCR activation can be organized into two categories: a dependence on OCT3 cationic channel-permeable ligands or the necessity of endocytic trafficking. Using CXC chemokine receptor 4 (CXCR4) as a model, we identified a third mechanism of intracellular GPCR signaling. We show that independent of membrane permeable ligands and endocytosis, upon stimulation, plasma membrane and internal pools of CXCR4 are post-translationally modified and collectively regulate *EGR1* transcription. We found that β-arrestin-1 (arrestin 2) is necessary to mediate communication between plasma membrane and internal pools of CXCR4. Notably, these observations may explain that while CXCR4 overexpression is highly correlated with cancer metastasis and mortality, plasma membrane localization is not. Together these data support a model were a small initial pool of plasma membrane-localized GPCRs are capable of activating internal receptor-dependent signaling events.

---

While extracellular inputs, cell membrane receptors, and resulting transcriptional programs are diverse, many receptor-signaling events converge to a reduced number of signaling hubs. Cellular mechanisms that mediate this process as well as strategies to control these actions remain outstanding questions. Over the last decade, we have learned that GPCR spatiotemporal signaling is one mechanism used by cells to translate diverse environmental information into actionable intracellular decisions while using seemingly redundant signaling cascades^1^. Extensive research has illustrated that GPCRs elicit distinct signaling events at different plasma membrane micro-domains as well as endocytic compartments that are important for cell physiology and disease pathogenesis^1–9^. These studies support a model where the location, in addition to magnitude, of a signaling event is important for cellular decision-making. Others have shown that GPCR site-specific post-translational modifications (PTMs) modulate adaptor protein recruitment, GPCR localization, and consequently receptor signaling events^10–12^. Together these observations motivated us to reexamine some confounding observations pertaining to the relationship of receptor localization, PTM, and signaling for CXCR4.

CXCR4 is a type 1 GPCR that regulates a variety of biological processes such as cell migration, embryogenesis, and immune cell homeostasis^10,13–16^. It is deregulated in 23 different cancers and overexpression is often correlated with metastasis and mortality^17–21^. However, surprisingly, plasma membrane expression is not correlated with metastasis^21^ and in some cancer tumor specimens as well as cell culture models, samples with poor CXCR4 plasma membrane localization remain responsive to CXCR4 agonist^22–26^. CXCR4 is activated by a highly receptor-specific 8 kDa chemokine, CXCL12^27–29^. Unlike β-adrenergic receptors which have been shown to be activated at intracellular compartments in an OCT3 cationic transporter-dependent mechanism^4,7^, endocytic-independent internalization of CXCL12 is unlikely due to its size. Given that receptor activation is dependent on ligand binding or transactivation by another receptor^30,31^, the aforementioned observations are confounding, as cells with low plasma membrane CXCR4 remain highly responsive to CXCL12. There are two potential explanations for this observation. Firstly, this could be due to spare receptors on the plasma membrane as it is well established that only a limited number of plasma membrane receptor contribute to signaling^32^. Alternatively, this could be due to activation of intracellular pools of receptors.

We began first by investigating the role of CXCR4 localization on receptor signaling and PTM. To do so, we needed a strategy to robustly detect CXCR4 PTM as well as a method to modulate receptor localization. We previously established the use of a monoclonal CXCR4 antibody (UMB2) as a robust tool to study CXCR4 PTM^33^. This commercially available antibody is raised against the C-terminus of the receptor, and upon CXCL12 stimulus quickly loses its ability to detect CXCR4 due to receptor PTM^33^. To attempt to identify the specific PTMs responsible, we treated lysates from WT and ubiquitination mutant receptor expressing cells with phosphatase. While phosphatase treatment ablated AKT S473 phosphorylation, no change in UMB2 detection was observed (Supplemental Fig. 1a, b). Since it has also been reported that CXCR4 is methylated at C-terminal arginine residues^34,35^, we tested whether CXCR4 methylation was responsible for the agonist-dependent loss in UMB2 detection. Similar to phosphatase treatment, protein methylation inhibition did not affect UMB2 detection (Supplemental Fig. 1c). Together these results suggest that the agonist-dependent reduction in UMB2 antibody detection is likely due to a combination of CXCR4 PTMs.

To manipulate receptor localization, we generated several mutant receptors that modulate the steady-state distribution of CXCR4 within retinal pigment epithelial (RPE) cells (Fig. 1a). We chose RPE cells to study CXCR4 overexpression because they do not have appreciable endogenous CXCR4 expression, are unresponsive to CXCL12, and were previously established as a cell culture model to study CXCR4 biology^33^. We found that by mutating C-terminal lysine residues to arginine (K3R), CXCR4 plasma membrane localization was reduced by more than 50% (Fig. 1b, c). While the CXCR4 K3R mutant has been previously used to study CXCR4 degradation^36,37^, we found that these mutations caused a drastic change in the spatial distribution of CXCR4. This was due to the unintended creation of an R-X-R motif, which has been shown to increase GPCR retention in the Golgi^38–40^. Indeed, mutating a single residue in the R-X-R motif (i.e., K3R/Q) restored receptor plasma membrane localization to near WT levels (Fig. 1b).

**Fig. 1:**
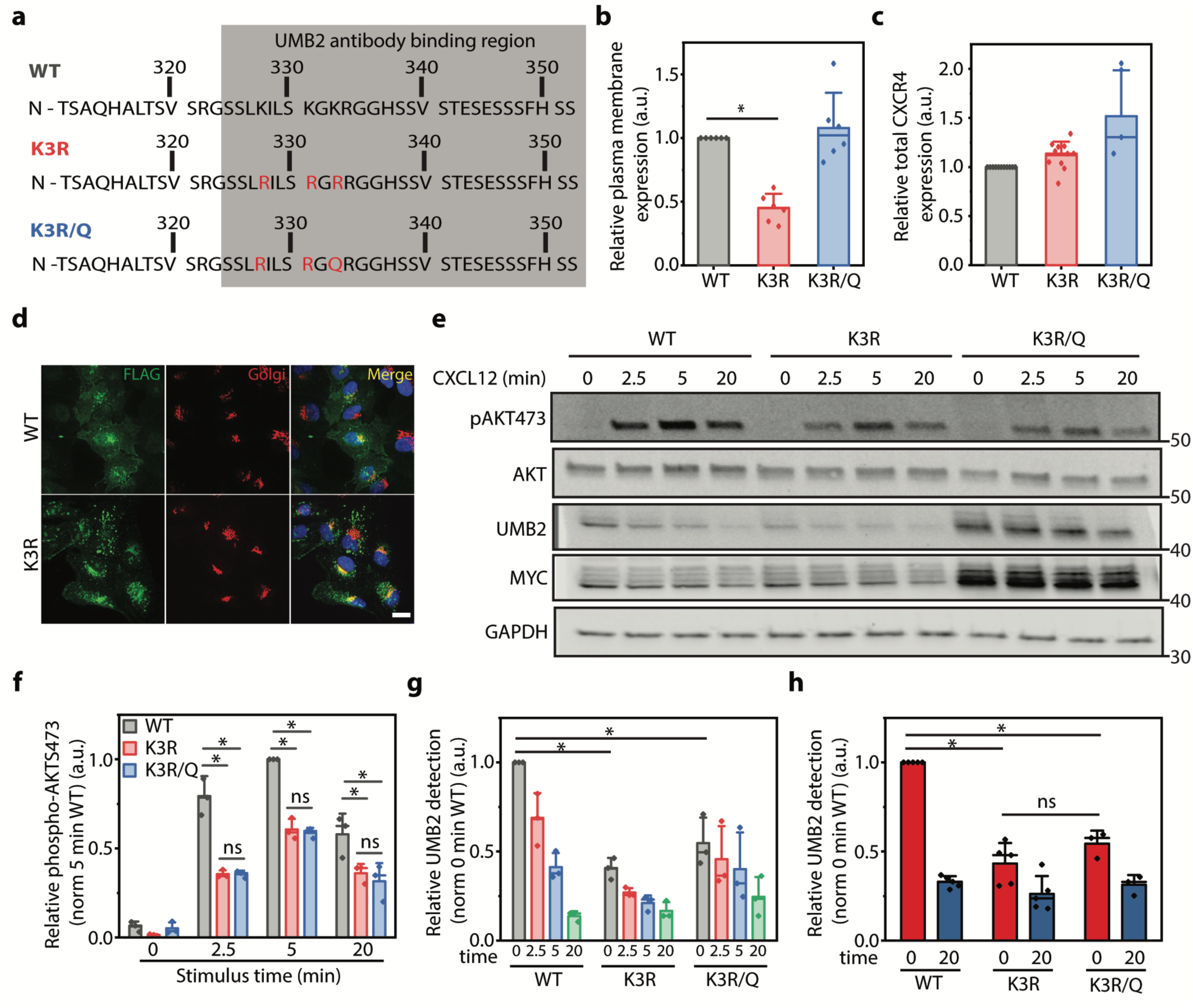
CXCL12-depedent AKT S473 phosphorylation and CXCR4 PTM are independent of CXCR4 localization. **a** Illustration of CXCR4 mutant receptor constructs. The gray box denotes the binding region of the UMB2 antibody that is sensitive to CXCR4 PTM. **b** Flow cytometry analysis of overexpressed WT and mutant receptor plasma membrane localization in RPE cells. Data was normalized to total receptor and WT plasma membrane expression. **c** Flow cytometry analysis of WT and mutant receptor total expression. Individual data points were normalized to WT CXCR4 expression. **d** Representative microscopy images illustrating the distribution of WT and K3R CXCR4 localization within RPE cells overexpressing each construct. CXCR4 was labeled with a FLAG antibody and the Golgi was detected using a GM130 antibody. Scale bar is 10 μm. Images were captured using 60x magnification on a spinning disk confocal microscope. **e** Representative western blot illustrating CXCL12-induced (12.5 nM) AKT S473 phosphorylation and CXCR4 PTM for WT and mutant receptors. Total CXCR4 was detected using a MYC antibody and unmodified CXCR4 by UMB2. **f** Western blot quantification of AKT S473 phosphorylation for WT and mutant CXCR4. Relative AKT phosphorylation was calculated by normalizing phospho-AKT to total AKT band intensity and secondly to the 5 min control time point. **g** Western blot quantification of CXCR4 PTM (i.e. UMB2 detection). A decrease in UMB2 detection is correlated to increased CXCR4 PTM. CXCR4 PTM was calculated by dividing the UMB2 intensity by the MYC intensity (total CXCR4) and secondly to the 0 min time point for the WT receptor. **h** Flow cytometry analysis of agonist-dependent WT and mutant CXCR4 PTM. Relative UMB2 detection was determined by dividing median UMB2 detection by total CXCR4 fluorescence and normalized to 0 min WT CXCR4. All experiments were conducted a minimum of 3 times in RPE cells overexpressing WT or mutant CXCR4. Individual data points from each experiment are plotted; mean, standard deviation (SD), median line. Statistical significance (*) denotes *p* < 0.05.

Interestingly, while total CXCR4 expression was unchanged by the K3R mutant, the K3R/Q mutant had slightly higher expression compared to WT receptor (Fig. 1c). In accordance with previous literature, K3R mutant receptors partially colocalized with a Golgi compartment marker (Fig. 1d)^38,39^. We also noticed a steady-state population of WT CXCR4 retained at the Golgi (Fig. 1d). Non-plasma membrane localized CXCR4 has been previously reported^41^ and could potentially be due to the presence of a K-X-K motif, which has also been implicated in Golgi protein retention^42–44^ or receptor overexpression. It is important to point out that while we observed partial CXCR4 colocalization with the Golgi, it is evident from our microscopy results that CXCR4 is present at other intracellular compartments as well (Fig. 1d).

Having established methods to modulate receptor localization and monitor PTM, we proceeded to investigate the role of CXCR4 plasma membrane localization on CXCL12-dependent AKT S473 phosphorylation. Since CXCL12 is not membrane permeable, we hypothesized that plasma membrane localization is essential for CXCR4 signaling and that reducing receptor plasma membrane expression would decrease CXCL12-dependent AKT phosphorylation. Compared to WT, both mutants had significantly reduced CXCL12-dependent AKT phosphorylation (Fig. 1e). This was expected as mutating biologically relevant residues may affect G protein coupling to CXCR4. However, surprisingly, there was no difference in AKT phosphorylation between high (K3R/Q) and low (K3R) plasma membrane localized mutant receptors (Fig. 1e, f). Additionally, while not explicitly investigated, earlier studies using the K3R mutant also reported that CXCL12-induced ERK1/2 phosphorylation was similar between WT and K3R mutant receptor expressing cells^37^. Since receptor PTM plays a role in mediating receptor signaling, we investigated the effect of receptor localization on agonist-induced CXCR4 PTM. Although UMB2 detection continued to decrease over a course of three hours (Supplemental Fig. 1d), here we focused on the UMB2 detection in the first 20 minutes post stimulus since we were interested in how early CXCR4 PTM regulates cell signaling. We hypothesized that receptor plasma membrane localization is essential for agonist-dependent PTM as agonist-induced receptor PTM is believed to require ligand binding. Surprisingly, irrespective of plasma membrane localization, mutant receptor PTMs were similar (Fig. 1e, g, h). Relative to total receptor expression, initial detection using the UMB2 antibody for both mutant receptors was also reduced (Fig. 1e, g, h). We believe this is because these mutations occur in the UMB2 antibody-binding region of the receptor (Fig. 1a). Alternatively, this could suggest a difference in steady-state mutant CXCR4 PTM.

Intrigued by the observation of a similar degree of CXCR4 PTM despite vastly different plasma membrane localization, we wondered whether internal (non-plasma membrane) pools of CXCR4 could be post-translationally modified in response to agonist stimulation and contribute to signaling. To investigate this, we examined the localization of CXCR4 PTM during receptor signaling. As previously observed^33^, upon CXCL12 addition UMB2 detection was drastically reduced at both plasma membrane and intracellular compartments, suggesting that both plasma membrane and internal pools of CXCR4 are post-translationally modified (Fig. 2a). To test this directly, we developed an assay to selectively isolate plasma membrane proteins from whole cell lysate. We used a membrane-impermeable promiscuous biotin molecule to selectively label and immunoprecipitate plasma membrane proteins with accessible extracellular domains (Fig. 2b). Receptor internalization was blocked throughout these experiments to keep plasma membrane and internal receptor pools distinct. We hypothesized that only plasma membrane receptors would be post-translationally modified, as internal pools of receptors are inaccessible to ligand and endocytosis of plasma membrane receptors is blocked. Surprisingly, we found that both surface and internal pools of receptors were post-translationally modified after ligand addition (Fig. 2c, d). To ensure that the labeling strategy was working as expected, we probed for GAPDH and biotinylated proteins and showed enrichment in expected localizations (Fig. 2c). To further examine intracellular CXCR4 PTM, we utilized blocking antibodies to effectively tune plasma membrane activity of endogenous CXCR4 in HeLa cells by varying the concentration of blocking antibodies (Fig. 2e). While the blocking antibody effectively reduced steady-state non-post-translationally modified CXCR4 (Fig. 2f), agonist addition had no effect on CXCR4 PTM after 20 min stimulus irrespective of CXCR4 plasma membrane expression level (Fig. 2g). Together these data support a model where internal pools of CXCR4 are post-translationally modified in response to CXCL12. Additionally, plasma membrane proteins are required for this process as removal of plasma membrane extracellular motifs using protease treatment completely ablated CXCL12-dependent CXCR4 PTM (Supplemental Fig. 2).

**Fig. 2:**
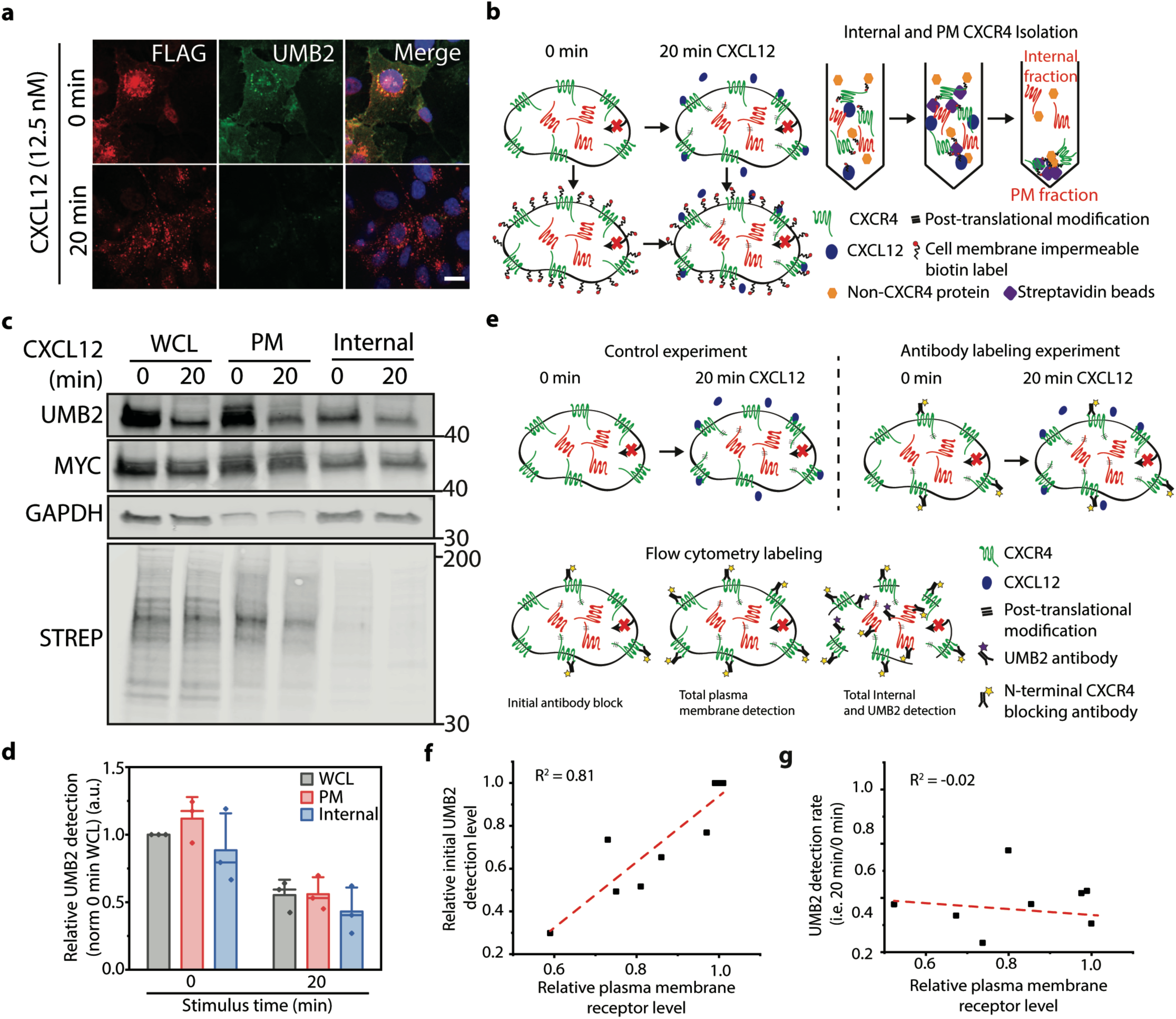
Plasma membrane and internal pools of CXCR4 are post-translationally modified upon receptor stimulus. **a** Representative microscopy images of total and non-post-translationally modified CXCR4 pre and 20 min post CXCL12 (12.5 nM) stimulation. Total CXCR4 is detected by FLAG antibody and non-PTM CXCR4 by UMB2. Images were captured using 60x magnification on a spinning disk confocal microscope. Scale bar is 10 μm. **b** Plasma membrane and internal CXCR4 isolation assay schematic. Receptor internalization was blocked using Dynasore (100 μM) throughout the experiment. At the completion of CXCL12 stimulation, plasma membrane proteins from control and CXCL12-treated samples were covalently labeled using promiscuous, membrane impermeable NHS-sulfo-biotin. Afterwards, plasma membrane proteins were isolated from whole cell lysate (WCL) by immunoprecipitation and WCL, plasma membrane, and internal pools of CXCR4 were analyzed for PTM by western blot. **c** Representative western blot showing CXCL12-dependent (12.5 nM) CXCR4 PTMs of WCL, plasma membrane, and internal CXCR4. STREP and GAPDH were used as experimental validation. **d** Quantification of CXCR4 PTMs at plasma membrane and internal locations. CXCR4 PTMs were calculated by dividing UMB2 detection (non-PTM CXCR4) by MYC intensity (total CXCR4) and normalized to the 0 min WCL sample. **e** Experimental schematic for antibody blocking experiment. Incubating cells with different concentrations of CXCR4 antibody reduced plasma membrane-localized, ligand-accessible, CXCR4. Afterwards, total plasma membrane, total CXCR4 and PTM CXCR4 were quantified by flow cytometry. Experiments were conducted in HeLa cells stimulated with or without 12.5 nM CXCL12. CXCR4 was blocked using various dilutions of a 12G5 allophycocyanin-conjugated CXCR4 antibody. Receptor internalization was blocked throughout all experiments using Dynasore (100 μM). **f** Initial antibody block reduced relative CXCR4 expression and basal UMB2 detection. **g** Relative UMB2 detection 20 min post CXCL12 stimulus plotted against relative CXCR4 plasma membrane expression. R^2^ values are shown for each experiment. All experiments were conducted in RPE cells overexpressing CXCR4 unless noted. A minimum of 3 independent replicates were conducted for all experiments and individual data points from each experiment are plotted; mean, SD, median line.

Our findings so far suggest that a signaling cascade may be responsible for intracellular communication between plasma membrane and internal receptor pools. Next, we focused on identifying the proteins responsible for agonist-dependent internal CXCR4 PTM. G proteins are master regulators for GPCR signaling and recent studies have revealed tight spatiotemporal regulation of G protein signaling events^6,7,45^. Upon ligand binding, G protein G_αi_ and G_βγ_ subunits are released from the GPCR due to guanidine exchange factor activity. G_βγ_ activates GPCR kinases (GRKs), which quickly phosphorylate the C-terminus of activated receptors leading to β-arrestin recruitment (Fig. 3a)^46^. Therefore, we hypothesized that G_βγ_ inhibition would reduce CXCL12-induced signaling and consequentially CXCR4 PTM. Indeed, pharmacological inhibition of G_βγ_ signaling significantly reduced both ERK1/2 and AKT phosphorylation (Fig. 3b-d). Additionally, G_βγ_ inhibition completely ablated CXCL12-dependent CXCR4 PTM (Fig. 3e, f).

**Fig. 3:**
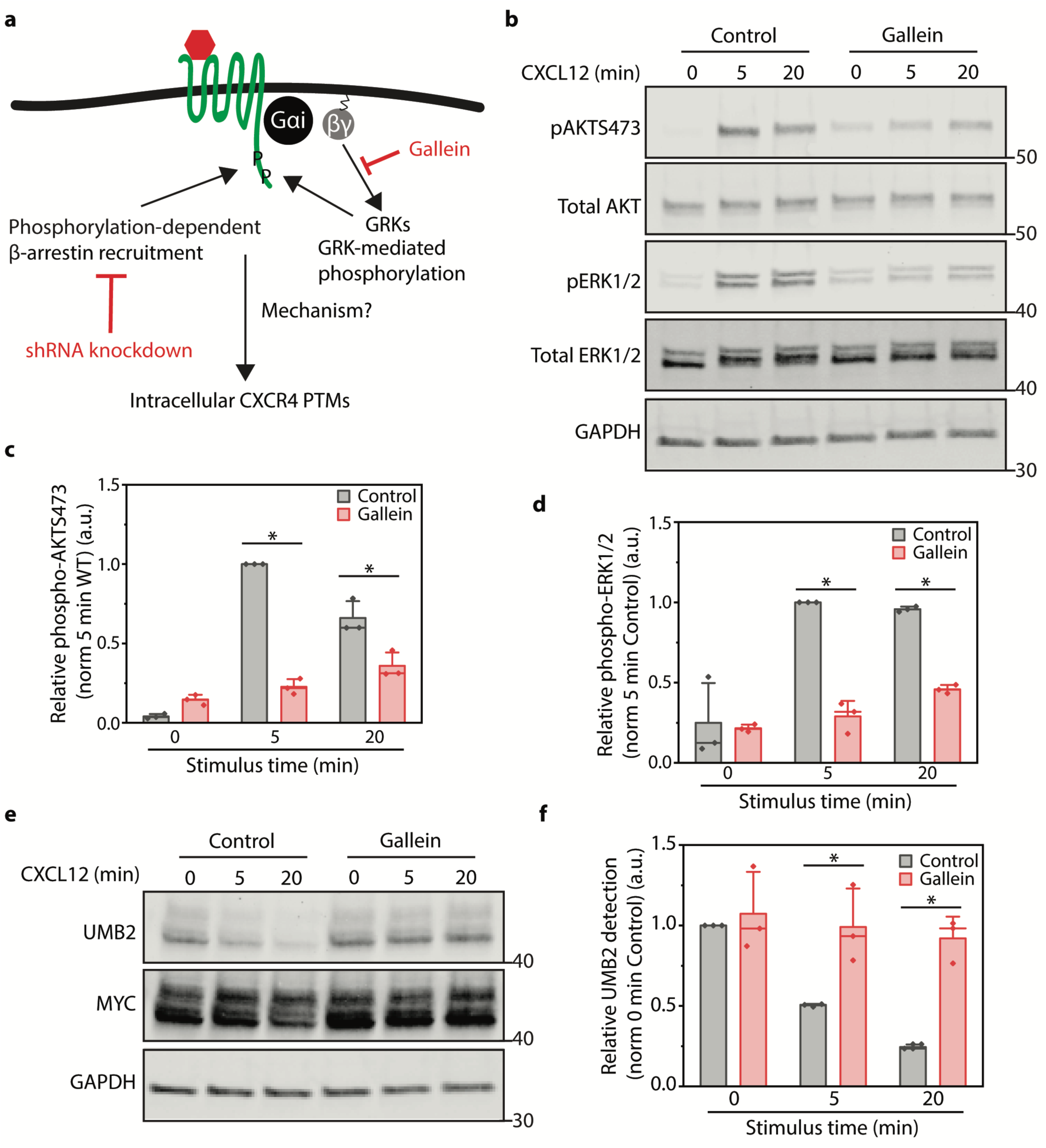
G_βγ_ signaling is essential for CXCR4 signaling and PTM. **a** Illustration of the current model of GPCR desensitization. Perturbations used to antagonize different components of the pathway are highlighted in red. **b** Representative western blot illustrating the effects of G_βγ_ inhibition (Gallein, 10 μM) treatment on CXCL12-induced AKT S473 and ERK1/2 phosphorylation. Cells were pretreated with Gallein for 30 min prior to and throughout each signaling time course. **c-d** Western Blot quantification of AKT S473 and ERK1/2 phosphorylation after G_βγ_ inhibition. Relative signaling protein phosphorylation was calculated by dividing the phosphorylated protein detection by total signaling protein detection and then normalized to the 5 min time point of the control sample. **e** Representative western blot illustrating the effect of G_βγ_ inhibition (Gallein 10 μM) on CXCR4 PTM. Cells were pretreated with Gallein for 30 min prior to and throughout the signaling time course. **f** Western blot quantification of CXCR4 UMB2 detection (i.e. PTM) upon G_βγ_ inhibition. CXCR4 PTM was calculated by dividing UMB2 detection (non-PTM CXCR4) by MYC intensity (total CXCR4) and normalized to the 0 min control sample. For all experiments a minimum of 3 independent replicates were performed. All experiments were conducted in RPE cells overexpressing WT CXCR4 and stimulated with 12.5 nM CXCL12 for the stated time course. Individual data points from each experiment are plotted; mean, SD, median line. Statistical significance (*) denotes *p* < 0.05.

Since G_βγ_ activation leads to GPCR phosphorylation and β-arrestin recruitment, we decided to investigate whether β-arrestins were involved in regulating internal CXCR4 PTM. β-arrestins have been previously implicated as potent messenger molecules. Coined signaling at a distance, work from the von Zastrow group proposed a new model for β-arrestin-dependent MAPK signaling in which β-arrestin-2, activated by stimulated GPCRs on the plasma membrane, traffics to nearby clathrin-coated structures to initiate localized MAPK signaling^8,47^. β-arrestin-1 and 2 are not equal, and significant research has revealed potential site-specific PTM and kinase phosphorylation-specific recruitment to CXCR4 as well as other GPCRs^11^. We hypothesized that β-arrestin 1 or 2 are important for communication between plasma membrane and internal CXCR4 (Fig. 3a). β-arrestin-1 knockdown led to a reduction in agonist-dependent CXCR4 PTMs while β-arrestin-2 knockdown had no effect (Fig. 4a). β-arrestin-1 knockdown did not affect ERK1/2 phosphorylation but led to a slight increase in AKT phosphorylation (Fig. 4b-e). This is potentially due to a failure to arrest G protein signaling. Intrigued by the potential new role of β-arrestin-1 in regulating the communication between plasma membrane and internal pools of receptors, we used the plasma membrane biotinylation assay to determine which CXCR4 population is regulated by β-arrestin-1. While β-arrestin-1 knockdown did not affect plasma membrane-localized CXCR4 PTM, internal CXCR4 PTM was reduced (Fig. 4f-h). Together these data support a mechanism by which G_βγ_ and β-arrestin-1 work together to regulate communication between plasma membrane and internal pools of CXCR4.

**Fig. 4:**
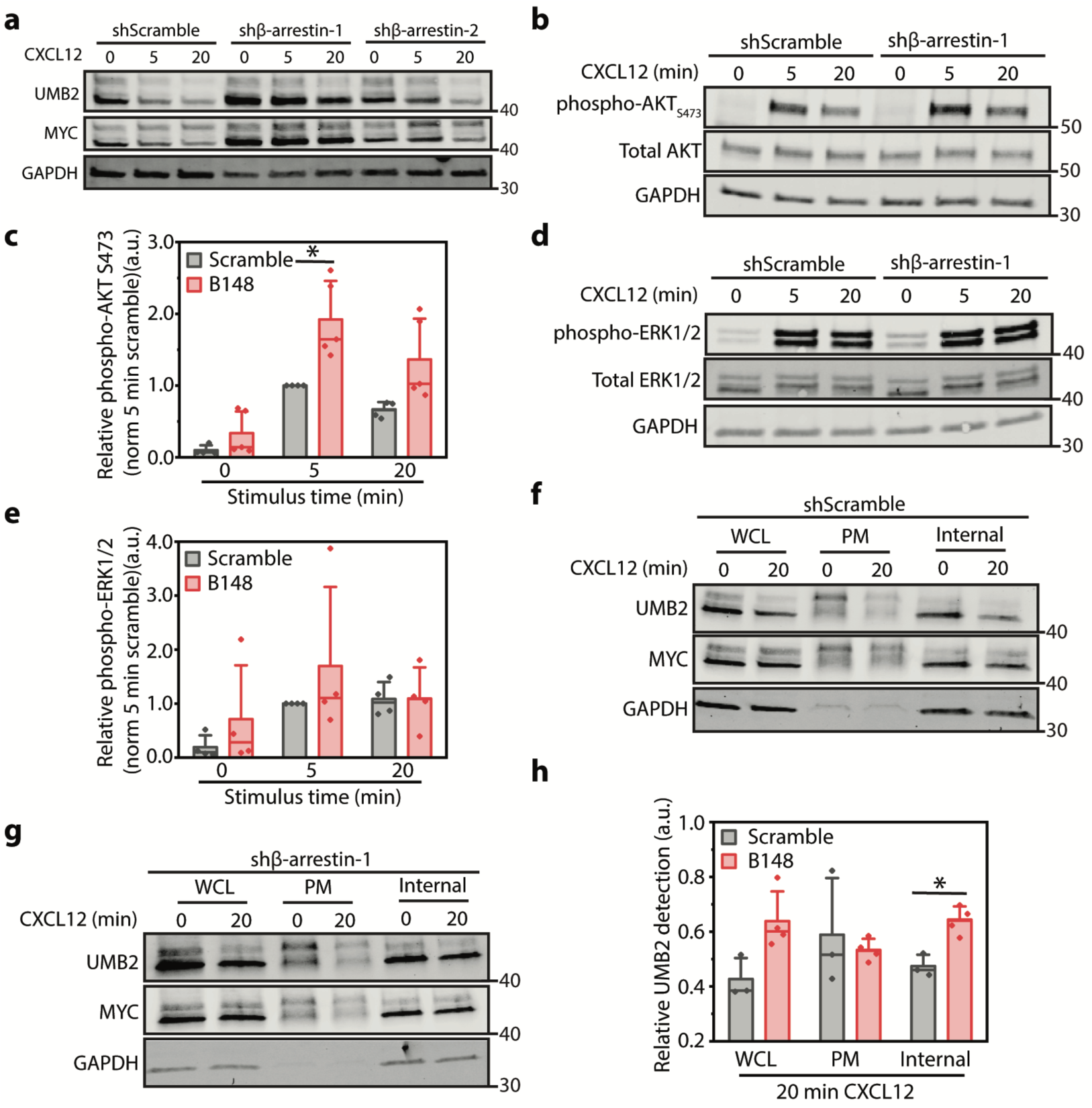
β-arrestin-1 regulates agonist-induced internal CXCR4 PTM. **a** Representative western blot illustrating the effect of β-arrestin-1 and β-arrestin-2 knockdown on CXCR4 PTM. Relative shRNA knockdown efficiency is shown in Supplemental Fig. 3a-c. **b** Representative western blot illustrating the effect of β-arrestin-1 knockdown on CXCL12-dependent AKT S473 phosphorylation. **c** Western blot quantification of CXCL12-dependent AKT S473 phosphorylation upon β-arrestin-1 knockdown. Data was normalized to phospho-AKT:total AKT and to 5 min normalized control shRNA sample. **d** Representative western blot illustrating the effect of β-arrestin-1 knockdown on CXCL12-dependent ERK1/2 phosphorylation. **e** Western blot quantification of CXCL12-dependent ERK1/2 phosphorylation upon β-arrestin-1 knockdown. Data was normalized to phospho-ERK1/2:total ERK1/2 and to 5 min normalized control shRNA sample **f-g** Representative western blots illustrating total, plasma membrane, and internal pools of CXCR4 PTM upon either scramble or β-arrestin-1 shRNA knockdown. **h** Quantification of CXCR4 PTM at plasma membrane and internal locations upon β-arrestin-1 knockdown. CXCR4 PTMs were calculated by dividing UMB2 detection (non-post-translationally modified CXCR4) by MYC intensity (total CXCR4) and normalized to the 0 min time point at each location. For all experiments a minimum of 3 independent replicates were performed. All experiments were conducted in RPE cells overexpressing WT CXCR4 and stimulated with 12.5 nM CXCL12 for the stated time course. β-arrestin-1 knockdown experiments were conducted using two validated shRNAs (Supplemental. Fig 3). Individual data points from each experiment are plotted; mean, SD, median line. Statistical significance (*) denotes *p* < 0.05.

Since non-plasma membrane CXCR4 overexpression has been associated with metastatic potential^22,26^, we wanted to investigate whether the internal pool of CXCR4 activated distinct signaling pathways compared to plasma membrane-localized receptors. GPCR signaling at intracellular compartments has become increasingly apparent and has been shown to activate different signaling cascades compared to plasma membrane-localized counterparts^1,3,48,49^. Recent work has shown that activation of G protein signaling at the Golgi and endosomes regulates PI4P hydrolysis and *PCK1* transcription respectively^7,49^. Therefore, we investigated whether activation of intracellular CXCR4 differentially activates downstream signaling compared to plasma membrane receptors. Rather than using mutant receptors that have preexisting signaling defects (Fig. 1), we decoupled the effects of CXCR4 localization from these mutations by removing a synthetic plasma membrane localization sequence commonly used to increase CXCR4 plasma membrane trafficking^36^. Consistent with our earlier findings, AKT and ERK1/2 phosphorylation as well as total CXCR4 PTM were not affected by modulating receptor localization (Supplemental Fig. 4). Since no overt defect in signaling was observed, we hypothesized that signal location is responsible for differential CXCL12-dependent transcription. To investigate this, we measured early growth response gene 1 (*EGR1)* transcript levels upon CXCL12 stimulus in RPE cells with high and low plasma membrane CXCR4 expression. *EGR1* transcription is downstream of the ERK1/2 pathway and has been shown to be induced by CXCL12^50,51^. As expected, WT RPE cells (not overexpressing CXCR4) were unresponsive to CXCL12 (Fig. 5a). However, compared to cells with high plasma membrane expression, cells with low plasma membrane CXCR4 expression had significantly increased CXCL12-induced *EGR1* transcript levels (Fig. 5a). This result is inconsistent with the spare receptor model. Furthermore, in agreement with previous work^6,7,49^, this suggests that while cells often use some of the same signaling machinery, the localization of a signaling event can lead to different cellular responses. Since β-arrestin-1 plays a role in activating internal CXCR4, we investigated whether inhibition of β-arrestin-1 decreased CXCL12-induced *EGR1* transcription. Indeed, β-arrestin-1 knockdown reduced agonist-induced *EGR1* transcript levels (Fig. 5b), providing additional evidence that intracellular pools of CXCR4 are physiologically relevant and that their function is dependent on β-arrestin-1.

**Fig. 5:**
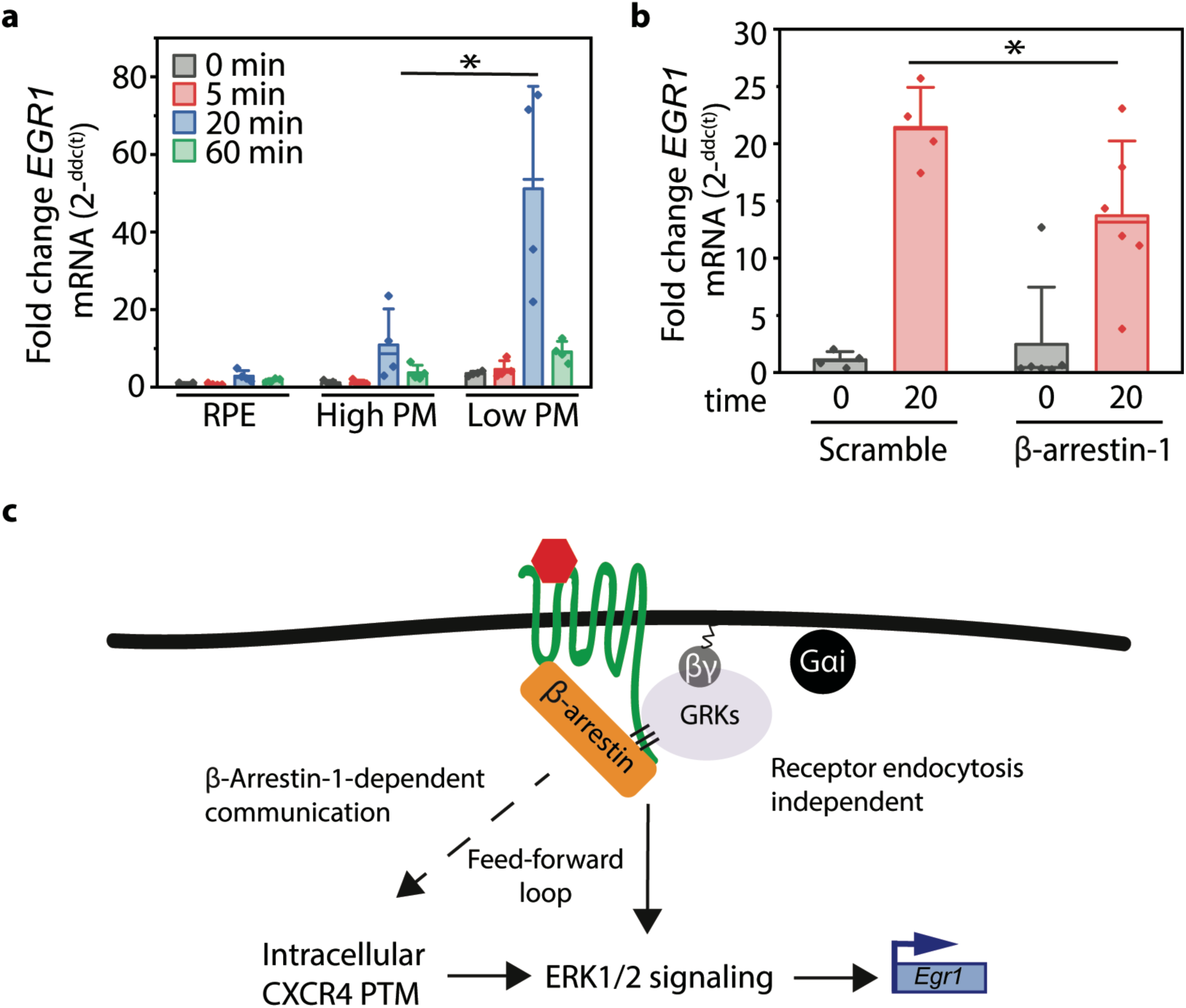
Intracellular pools of CXCR4 are primarily responsible for *EGR1* transcription. **a** qPCR analysis of *EGR1* transcription in WT RPE or RPE cells overexpressing high or low plasma membrane localized CXCR4. *EGR1* transcript levels were calculated using the ΔΔCT method normalized to GAPDH and 0 min high plasma membrane CXCR4. **b** β-arrestin-1 knockdown reduces CXCL12-dependent *EGR1* transcript levels in RPE cells overexpressing CXCR4. *EGR1* transcript levels were calculated using the ΔΔCT method. **c** Schematic summarizing a potential model for communication between plasma membrane and internal GPCR pools. All experiments were conducted in RPE cells overexpressing WT CXCR4 and stimulated with 12.5 nM CXCL12 for the stated time course unless noted. β-arrestin-1 knockdown experiments were conducted using two validated shRNAs (Supplemental Fig. 3). Individual data points from each experiment are plotted; mean, SD, median line. Statistical significance (*) denotes *p* < 0.05.

This work has revealed a new element of GPCR signaling whereby plasma membrane and internal pools of CXCR4 communicate to regulate cell signaling. These observations are distinct from previous work investigating intracellular GPCR signaling, where receptor internalization or OCT3 channel-permeable ligands were required^2–4,7,52,53^. CXCR4 PTM is dependent on G_βγ_ activation and β-arrestin-1 plays a specific role in regulating intracellular CXCR4 PTM. This work expands upon the β-arrestin signaling at a distance concept and supports a model where β-arrestins are not only able signal at a distance at the plasma membrane but also regulate communication between plasma membrane and internal GPCR populations to influence agonist-dependent transcriptional programs. A model for communication between plasma membrane and intracellular pools of CXCR4 and its ramification on signaling is summarized in Fig. 5c. There are several potential mechanisms for how this communication may occur that warrant additional research.

Many new questions regarding the molecular mechanism and physiological relevance of plasma membrane and intracellular CXCR4 activation remain unanswered. A limitation of our surface biotinylation approach to study intracellular CXCR4 PTMs is that it is unable parse out which subpopulation(s) of CXCR4 are being post-translationally modified. While agonist-induced internal CXCR4 PTM appear to partially occur at the Golgi (Fig. 2a), CXCR4 may also be post-translationally modified at other intracellular compartments. Understanding which intracellular CXCR4 populations are post-translationally modified is important for understanding which receptor pool is responsible for CXCL12-induced *EGR1* transcription. Furthermore, neither β-arrestin or G_βγ_ are believed to actively post-translationally modify proteins. However, G_βγ_ is a potent kinase activator therefore identification of the kinase or other protein machinery responsible for internal CXCR4 PTMs is necessary. Interestingly, G_βγ_ has been shown to traffick to intracellular compartments including the Golgi after agonist activation, independent of receptor endocytosis^54,55^. Understanding the specific PTMs of plasma membrane and internal CXCR4 populations could also provide important insights pertaining to the function and fate of activated intracellular receptors. It is possible that this mechanism may regulate receptor trafficking to the plasma membrane, effectively providing cells with a short-term memory of prior signaling events.

While many questions remain, the data presented expand the role of G_βγ_ and β-arrestins in regulating GPCRs and support a new model of GPCR signaling whereby plasma membrane and internal pools of receptors communicate to collectively determine a cellular response. These observations may resolve the paradox that while CXCR4 overexpression is associated with metastatic potential, plasma membrane localization is not^22,23^. Additionally, this work supports a growing amount of evidence supporting that targeting specifically intracellular GPCR populations or the downstream signaling cascades activated by these pools might lead to improved therapeutic strategies for treating cancer, cardiovascular disease, and pain management^6,7,40^.

## Acknowledgements

MSD thanks support from the National Science Foundation Graduate Research Fellowship and the University of Michigan Rackham Pre-doctoral Fellowship. The work is supported in part by Pardee Foundation and a gift from Kendall and Susan Warren. MSD would also like to thank Greg Thurber for allowing us to use his LiCor CLX imager for preliminary work in this study. MSD would like to thank Manoj Puthenveedu for discussion and Wylie Stroberg for data interpretation feedback.

## Methods and Reagents

### Equipment

LiCor Odessey CLX & SA Imagers

Azure Sapphire 4 laser Imager

BioRad RT-qPCR ThermoCycler

### Cell culture

HeLa cells were originally obtained from ATCC. HeLa cells were cultured in DMEM media (Corning) supplemented with 10% FBS (Corning). Retinal pigment epithelial (RPE) cells were a gift from Dr. Sandra Schmid at UT Southwestern. All stable cell lines were directly derived from this RPE line. RPE cell lines were cultured in DMEM/F12 media (Corning) supplemented with HEPES, glutamate and 10% FBS (Corning). HEK293T cells were obtained from ATCC and grown in DMEM (Corning) media supplemented with 10% FBS.

### DNA constructs and stable cell lines

WT CXCR4 was generated as previously described^33^. K3R and K3R/Q mutant receptors were generated by PCR mutagenesis of WT CXCR4 in the pLVX plasmid using the NEB Quick-change mutagenesis kit. The low plasma membrane CXCR4 construct was generated by PCR amplification (excluding the 5’ plasma membrane HA localization peptide) and restriction enzyme cloning using the *BsrRI* and *EcoRI* restriction enzymes. All CXCR4 constructs had an N-terminal FLAG tag and C-terminal MYC tag for easy antibody detection. Stable cell lines expressing WT and mutant CXCR4 receptors were generated by lentiviral transduction. Lentiviruses (shRNA and CXCR4 constructs for stable cell lines) were generated by co-transfecting HEK293T cells with the pLVX transfer plasmid, psPAX2, and pMD2.G lentiviral envelope and packaging plasmids. To generate stable cell lines, supernatant media containing mature lentiviral particles was collected 4 days post transfection and added to RPE cells, and cells stably expressing the constructs were generated via puromycin selection (3 µg/mL). All transfections were conducted using Lipofectamine 2000 (Life technologies).

### Flow cytometry experiments

Flow cytometry experiments for plasma membrane receptor labeling were conducted as previously described^33^. For intracellular staining, cells were first disassociated using 50 µM EDTA in Ca^2+^-free PBS and fix for 10 min in 4% paraformaldehyde at room temperature. Afterwards, cells were permeabilized using 0.2% Triton-X 100 for 10 min at room temperature. Intracellular targets were labeled with primary antibodies for 1 hr at room temperature after which cells were washed with PBS and incubated with secondary antibodies for 1 hr at room temperature – see Table 1 for antibody specifics. Afterwards, cells were washed 1x with PBS and 25,000 events were analyzed by the Guava EasyCyte flow cytometer for each experimental condition. When co-staining, compensation was conducted post experiment using controls with either 488 or 640 fluorescence alone. After fluorescence compensation, the median fluorescence was calculated for each channel and sample as well as for no stain and RPE WT controls (not expressing CXCR4). As previously described, median control sample fluorescence was subtracted from each sample and data was normalized and plotted as described in each figure legend^33^.

**Table 1.**

| **Inhibitors** | **PN** | **Supplier** | **Working Concentration** |
| --- | --- | --- | --- |
| Gallein | 3090/50 | R&D Systems | 10 μM |
| Dynasore | 324410 | Sigma | 100 μM |
| MTA | D5011 | Thermo Fisher | 200 μM |
| **Antibodies** |  |  |  |
| ms-FLAG-647 | A01811-100 | Genscript | 1:1000 (FC/IF) |
| rb-MYC | A190-105A | Bethyl Laboratories | 1:5000 (WB) |
| rb-CXCR4 (UMB2) | Ab124824 | Abcam | 1:2000 (WB),<br>1:1000 (FC/IF) |
| Rb-phospho-ERK1/2 | 4370S | Cell Signaling Technologies | 1:2000 (WB) |
| ms-total-ERK1/2 | 4696S | Cell Signaling Technologies | 1:1000 (WB) |
| rb-phospho-AKT S473 | 4060S | Cell Signaling Technologies | 1:2000 (WB) |
| rb-total AKT | C67E7 | Cell Signaling Technologies | 1:1000 (WB) |
| rb-GM130 | 12480S | Cell Signaling Technologies | 1:1000 (IF) |
| ms-GAPDH | sc-47724 | Santa Cruz Biotechnology | 1:1000 (WB) |
| STREP-568 | S11226 | Thermo Fisher | 1:5000 (WB) |
| ms-CXCR4-APC (12G5) | FAB170A | R&D Systems | 1:200 – 1:2000 (FC) |
| rb-β-Arrestin-1 | 30036S | Cell Signaling Technologies | 1:1000 (WB) |
| gt-anti-rb Dylight 800 | SA5-35571 | Thermo Fisher | 1:5000 (WB) |
| gt-anti-ms Dylight 680 | 35518 | Thermo Fisher | 1:5000 (WB) |
| gt-anti-rb-488-AlexaFluor-Plus | A32731 | Thermo Fisher | 1:1000 (FC/IF) |
| Gt-anti-rb-555 | 84541 | Thermo Fisher | 1:1000 (IF) |
| Gt-anti-rb-Alexafluor-Plus | A32733 | Thermo Fisher | 1:1000 (IF) |
| Gt-anti-ms-488 | A10680 | Thermo Fisher | 1:1000 (IF) |
| Gt-anti-ms-647 | A21235 | Thermo Fisher | 1:1000 (IF) |
| **Biologics** |  |  |  |
| CXCL12 | 350-NS-050 | R&D Systems | 12.5 nM |
| Pronase | 10165921001 | Sigma | 0.1% Solution |
| Lambda Phosphatase | sc-200312A | Santa Cruz Biotechnology | Manual instructions |
| **Others** |  |  |  |
| Streptavidin agarose | 20361 | Thermo Fisher |  |
| DAPI | D9542 | Sigma | 1µg /mL |
| NewBlot PVDF Stripping buffer | 928-40032 | LiCor | Manual instructions |
| PVDF 0.22 µm membranes | IB401001 | Thermo Fisher | NA |
| NHS-Sulfo-LC-biotin | 21335 | Thermo Fisher | 1 mg/mL |
| ITAQ Universal SYBR Green | 1725121 | BioRad | Manual instructions |
| iScript cDNA Synthesis Kit | 1706891 | RioRad | Manual instructions |
| Quick-RNA miniprep | R1054 | Zymo Research | Manual instructions |

### Cell signaling, shRNA and inhibitor experiments

Cells were seeded in 12 well plates 24 hr prior to each signaling experiment achieving 70-80% confluence at the experimentation time. Cells were serum-starved in DMEM/F12 media without FBS for 4 hr prior to each signaling experiment. For inhibitor experiments (Gallein, Dynasore) cells were pretreated for 30 min with the respective inhibitors and throughout the signaling experiment. For shRNA experiments, cells were transduced with either scramble or β-arrestin-1 or 2 shRNA (Table 2) for 3 days. shRNA lentiviral particles were generated as described above. Afterwards cells were stimulated with 12.5 nM CXCL12 (R&D Systems) for the labeled time course. Samples were washed with PBS 1x and lysed using RIPA buffer supplemented with protease inhibitors (EDTA-free Peirce protease inhibitor cocktail) and phosphatase inhibitors (HALT Phosphatase Inhibitor). For Lambda Phosphatase experiments, phosphatase inhibitors were excluded in the lysis buffer. After incubating cells with lysis buffer for 10 min on ice, lysates were collected and centrifuged at 16,000 *g* for 45 min at 4 _°_C. Afterwards lysates were immediately stored at −20 _°_C or processed for immunoprecipitation or western blotting.

**Table 2.**
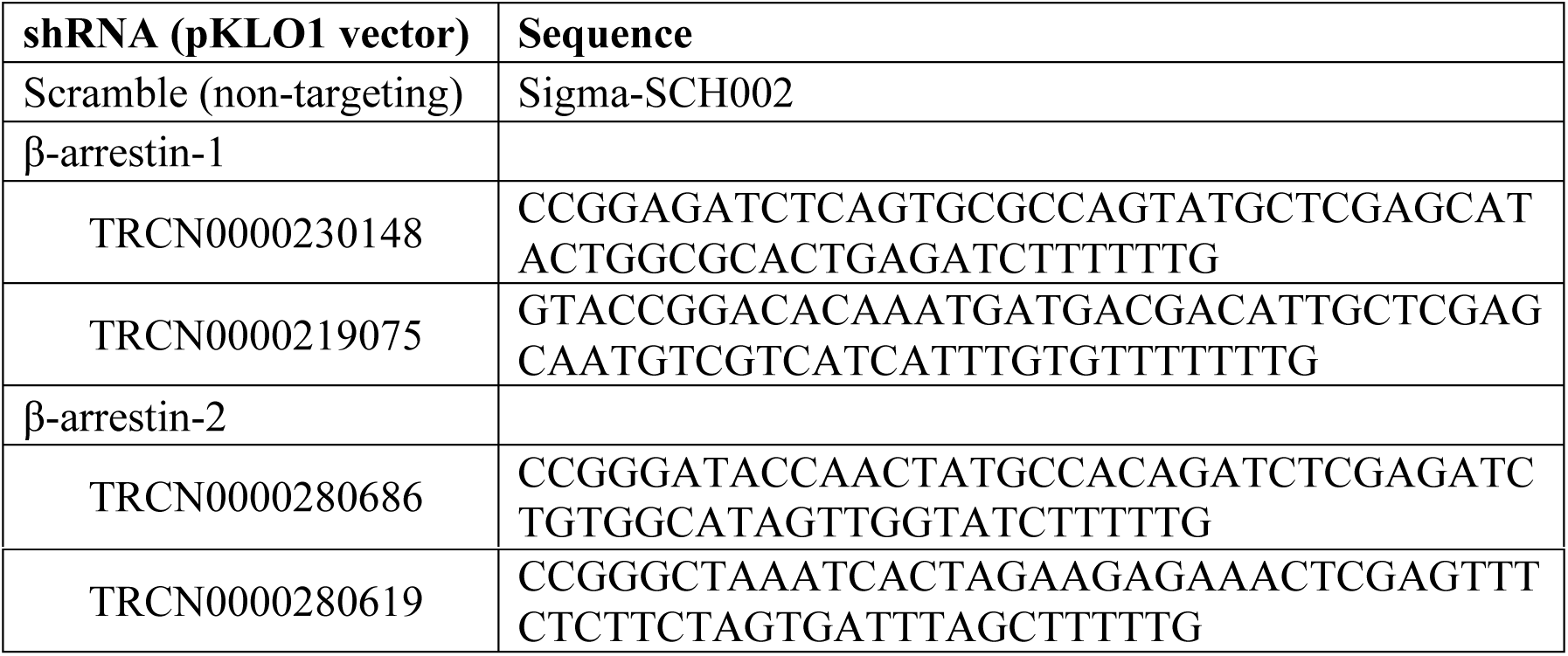

### Immunoprecipitation

For plasma membrane biotinylation experiments, biotinylated plasma membrane proteins were isolated from WCL using high capacity streptavidin agarose beads (Table 1). Approximately 35 µl of bead slurry was added to 350 µl WCL and incubated overnight (∼ 18 hr) rotating at 4 °C. Afterwards, samples were pelleted (centrifuged for 3 min at 2,000 *g*) and internal (non-plasma membrane) proteins collected by removing the supernatant. To prevent potential biotinylated protein contamination, only 200 µl of supernatant was removed. Afterwards, beads were washed 3 times with RIPA buffer containing protease inhibitors. After the final wash, all buffer was removed.

### Western blotting and data analysis

Prior to western blotting, samples were incubated with Laminelli buffer supplemented with β-mercaptoethanol (loading buffer). For surface biotinylation samples, β-mercaptoethanol concentration was increased 2-fold and samples were incubated at room temperature in the loading buffer for 30 min prior to western blotting to denature proteins from beads. Samples were run on SDS-PAGE 4-20% BioRad gels (15 well/15 µl or 10 well/50 µl gels). For all signaling experiments, 12.5 µl of lysate was loaded while for surface biotinylation assays, 35 µl of lysate was loaded. SDS-PAGE gels were run at constant 140 V for approximately 60 min. Afterwards, proteins were transferred to PVDF membranes using the iBlot transfer systems (mixed range proteins 7 min setting) and membrane incubated in blocking solution (1% BSA in TBST) rocking for 1 hr at room temperature. Afterwards, blots were incubated with their respective antibodies (Table 1) overnight at 4°C. Prior to secondary labeling, blots were wash 3x for 5 min per wash with TBST. Blots were then incubated with the corresponding secondary antibody (Table 1) for 1 hr at room temperature. Blots were then washed with TBST as described above. Western blots were dried and imaged using a LiCor Odessey SA, LiCor CLX, or Azure Biosystems Sapphire System. Data was analyzed using the LiCor image studio software to calculate band intensity as previously described^33^. Specific normalization procedures for each experiment are described in the respective figure legends. All statistics were calculated using two tailed t-tests.

### RT-qPCR experiments

Cells were seeded in 6 well plates 24 hr prior to each signaling experiment achieving 70-80% confluence at the experimentation time. Cells were serum-starved in DMEM/F12 media for 4 hr prior to each signaling experiment. Afterwards, cells were stimulated with 12.5 nM CXCL12 (R&D Systems) for the respective time courses shown in the figure legends. RNA was extracted using the Zymogen RNA extraction kit (R1054) and 1 µM cDNA was synthesized using the iScript synthesis kit (BioRad). qPCR assays were conducted using SYBR Green (BioRad) per BioRad protocol instructions using 12.5 ng of cDNA for each well. Samples were run in duplicate and primers used in this study are shown in Table 3. Samples were run on the BioRad CFX thermocycler and data was quantified using the ΔΔCT method as previously described^33^.

**Table 3.**
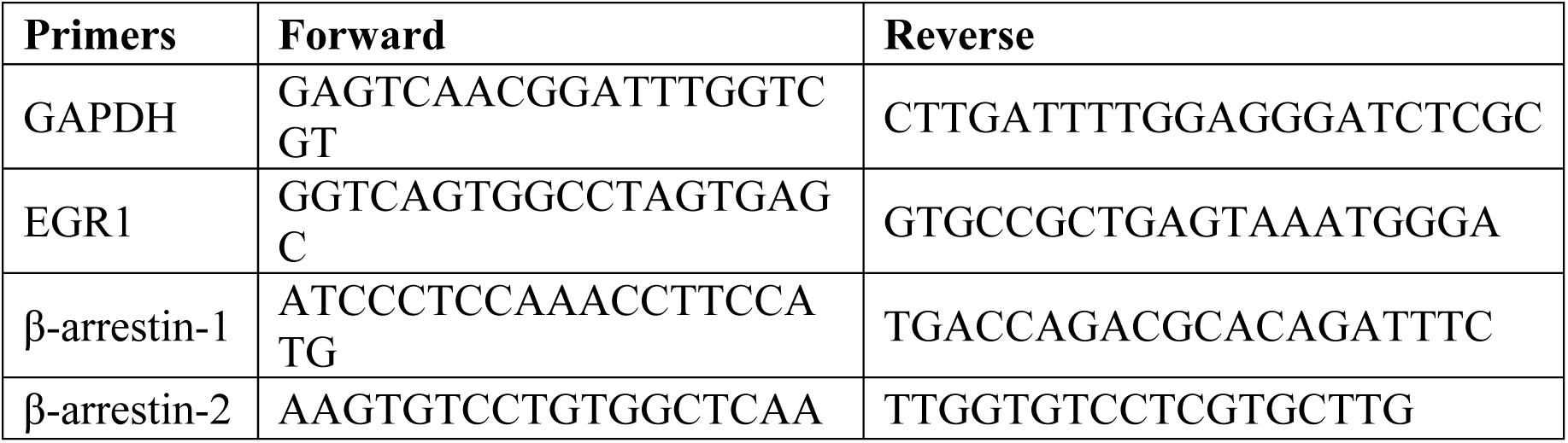

### Immunofluorescence Assays

Cells were seeded in 6-well plates on glass coverslips 24 hr prior to each experiment and serum-starved for 4 hr as described above. Cells were stimulated as specified in each figure legend and immediately washed with PBS and fixed in 4% paraformaldehyde for 10 min at room temperature. Cells were permeabilized for 10 min with 0.2% Triton-X100 diluted in PBS and subsequently blocked with 2.5% BSA diluted in PBS (blocking solution) for 1 hr. Cells were incubated with primary antibody diluted in blocking solution and incubated overnight at 4 _°_C (Table 1). Slides were wash 3x for 5 min each with PBS and incubated with secondary antibodies diluted in blocking solution for 1 hr at room temperature (Table 1). Cells were washed with PBS 3x, 5 min per wash and incubated with DAPI (Table 1) diluted in PBS for 10 min at room temperature. Afterwards, cells were washed with PBS and mounted onto glass slides using Fluoromount G. Slides were imaged by spinning disk confocal microscopy as specified in the figure legends. Different experimental samples were imaged using the same imaging settings each day.

