## Supplemental Figures for "β-arrestin mediates communication between plasma membrane and intracellular GPCRs to regulate signaling"

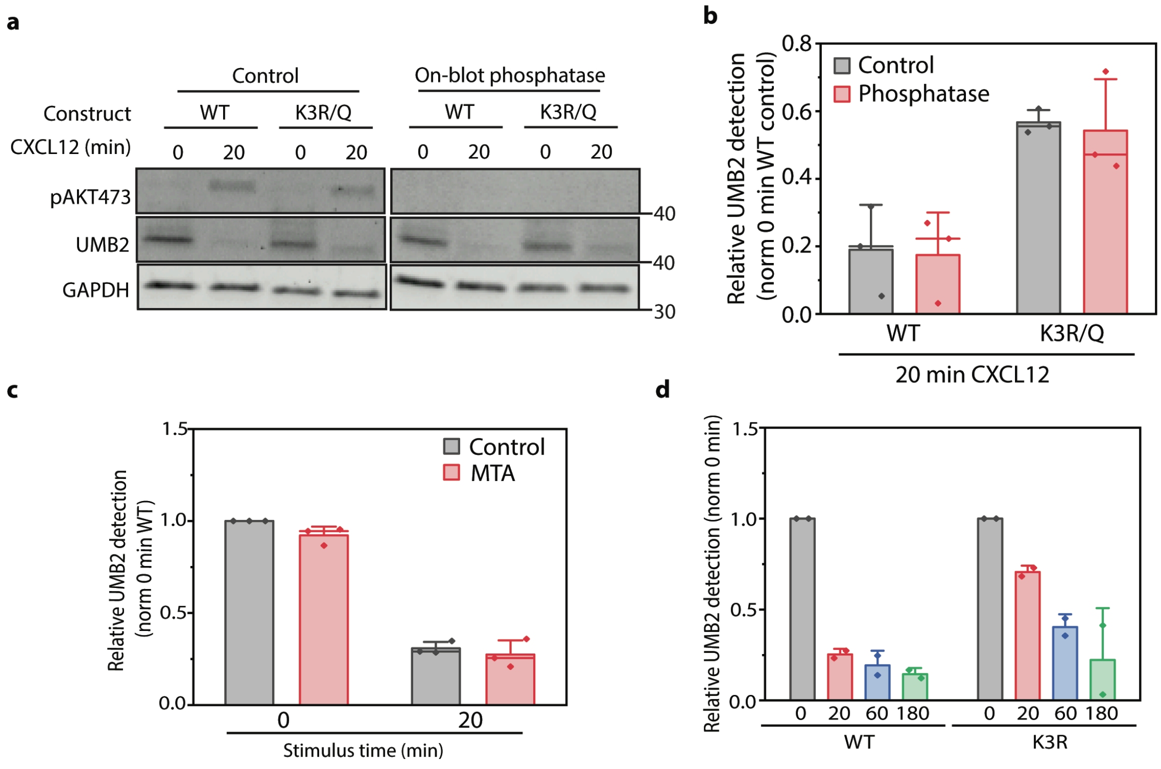


**Supplemental Fig. 1:** UMB2 antibody characterization. **a** Representative western blot of on-blot phosphatase experiment with WT and K3R/Q mutant CXCR4 constructs. The K3R/Q mutant receptor is unable to be ubiquitinated at the canonical C-terminal CXCR4 ubiquitination sites. The on-blot phosphatase blot was incubated with Lambda Phosphatase to remove phosphate groups from phosphorylated protein residues. Phospho-AKT S473 probing was included as a phosphatase treatment positive control. **b** Quantification of on-blot phosphatase western blotting experiments. Data was normalized by dividing the UMB2 detection at 20 min by the 0 min time point for either the WT or K3R/Q receptor. **c** Flow cytometry analysis of arginine protein methylation activity on CXCR4 UMB2 detection. As illustrated, cells were treated with methylthioadenosine (MTA 200 µM), an arginine protein methyltransferase inhibitor. UMB2 detection was normalized to total CXCR4 fluorescence and secondly to 0 min control detection. **d** Flow cytometry analysis of UMB2 antibody detection for an extended CXCL12-stimulus time course using WT or K3R mutant CXCR4. UMB2 detection was normalized to total CXCR4 fluorescence and secondly to 0 min detection for either WT or K3R mutant receptor. All experiments were conducted in RPE cells overexpressing WT CXCR4 and stimulated with 12.5 nM CXCL12 for the stated time course. Individual data points from each experiment are plotted; mean, SD, median line.

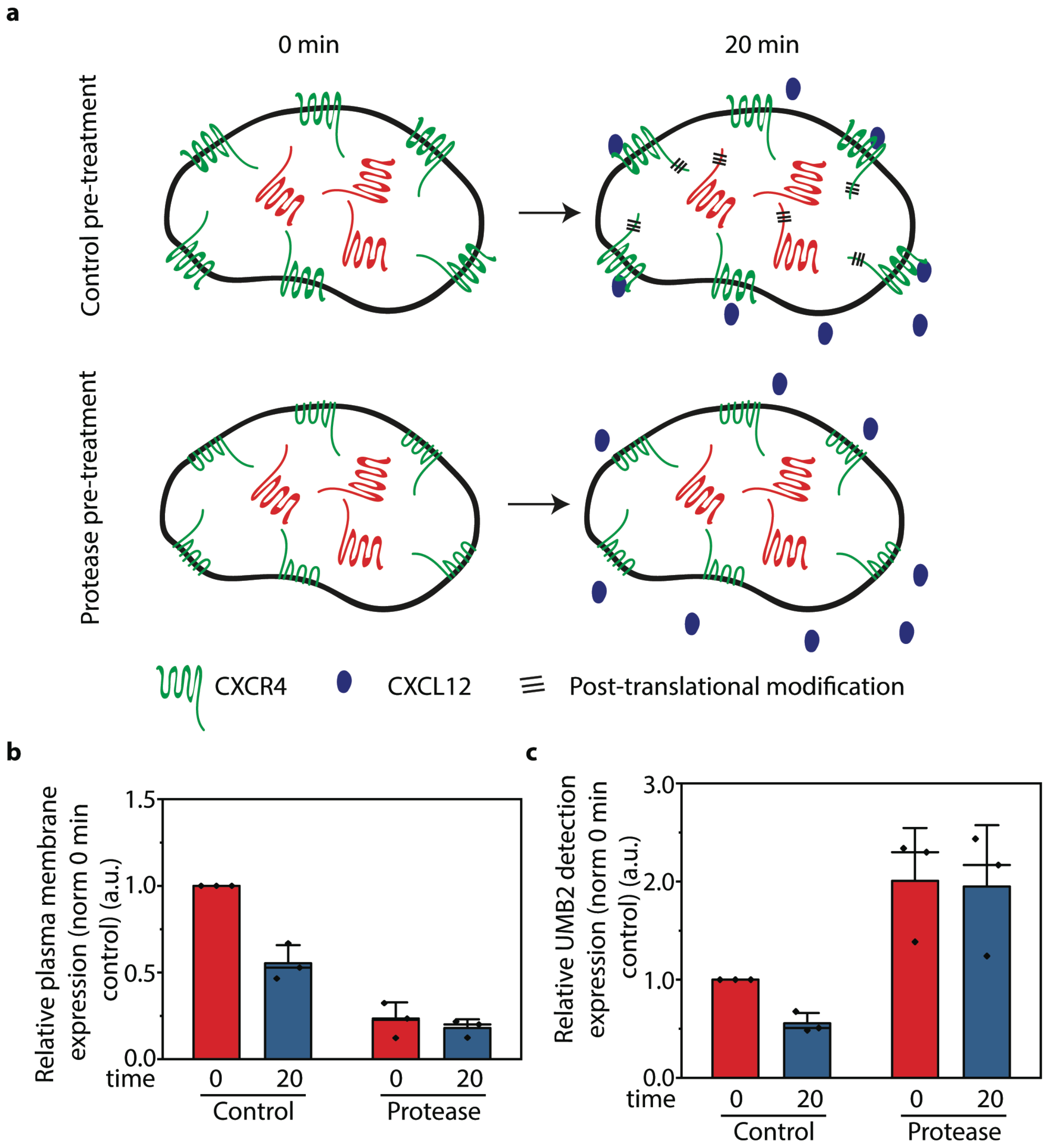


**Supplemental Fig. 2:** Functional plasma membrane proteins are required for CXCR4 PTM. **a** Experimental design schematic for the protease cleavage assay. Cells were pretreated with a protease (pronase 0.11% diluted in PBS) to remove extracellular motifs of transmembrane plasma membrane proteins. Afterwards, cells were stimulated with CXCL12 and plasma membrane CXCR4 and UMB2 detection was measured by flow cytometry. **b** Flow cytometry analysis of protease treatment on functional CXCR4 plasma membrane localization. Plasma membrane expression was measured using a FLAG antibody that detected an N-terminal FLAG-tagged CXCR4 and normalized to total CXCR4 expression. Data was normalized to the 0 min control sample. **c** Flow cytometry analysis of protease treatment on CXCR4 UMB2 detection. Relative UMB2 detection was calculated by dividing UMB2 signal by total CXCR4 signal (FLAG antibody) and then normalized to the 0 min control sample. An increase in UMB2 detection in the protease treatment sample is expected as the detectable total CXCR4 population is reduced due to protease cleavage of the N-terminal CXCR4 FLAG tag. All experiments were conducted in RPE cells overexpressing WT CXCR4 and stimulated with 12.5 nM CXCL12 for the stated time course. Individual data points from each experiment are plotted; mean, SD, median line.

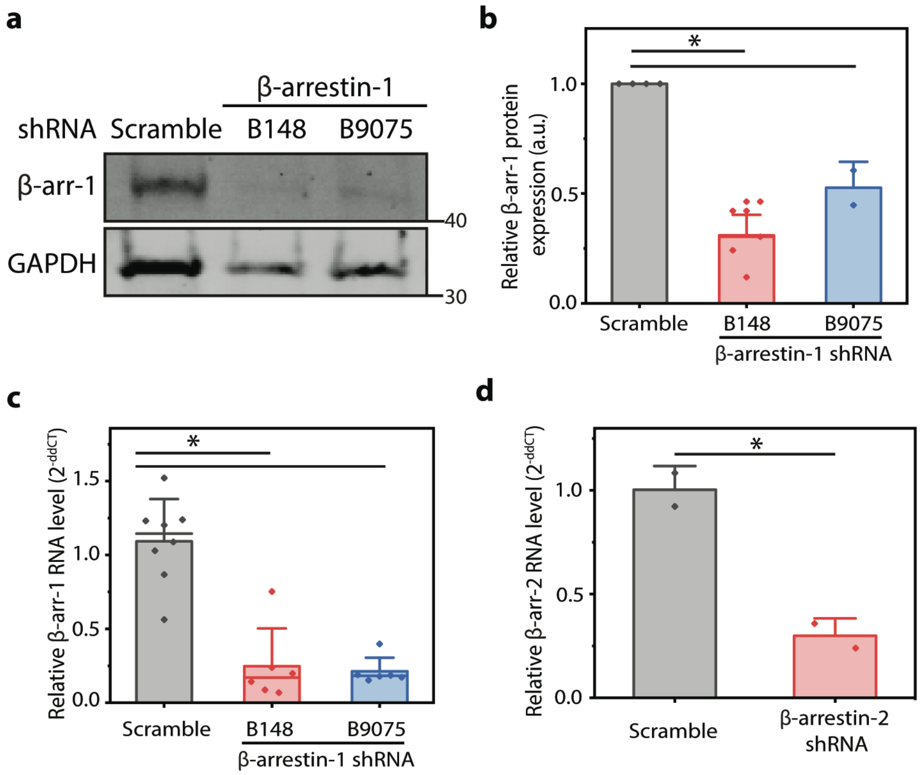


**Supplemental Fig. 3:** β-arrestin-1 knockdown confirmation. **a** Representative western blot illustrating relative β-arrestin-1 knockdown with two different shRNAs. **b** Quantification of β-arrestin-1 knockdown efficiency by western blot. Protein knockdown was calculated by dividing normalized β-arrestin-1 detection with β-arrestin-1 shRNA with scramble shRNA. **c-d** Relative β-arrestin-1 and β-arrestin-2 transcript levels were calculated using the ΔΔct method normalized to GAPDH and the scramble shRNA control. All experiments were conducted in RPE cells overexpressing WT CXCR4 and stimulated with 12.5 nM CXCL12 for the stated time course. Individual data points from each experiment are plotted; mean, SD, median line. Statistical significance (*) denotes $p < 0.05$.

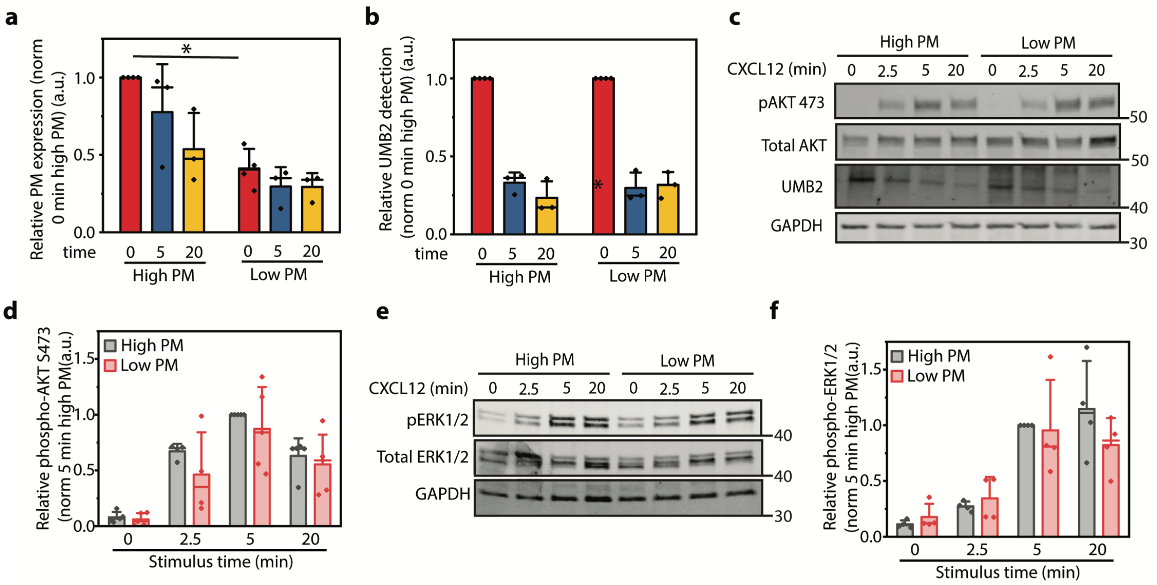


**Supplemental Fig. 4:** Modulation of CXCR4 plasma membrane localization does not affect CXCL12-dependent AKT S473 and ERK1/2 phosphorylation or CXCR4 PTM. **a** Flow cytometry analysis of CXCR4 plasma membrane expression of high and low plasma membrane (PM) expressing CXCR4 constructs. Plasma membrane expression was normalized to total CXCR4 expression and secondly to the 0 min high plasma membrane CXCR4 sample. **b** Flow cytometry analysis of high and low plasma membrane expressing WT CXCR4 constructs. UMB2 detection was normalized to total CXCR4 expression and secondly to the 0 min control samples. **c** Representative western blot illustrating CXCL12-induced AKT S473 phosphorylation and UMB2 detection in high and low plasma membrane expressing CXCR4 cells. **d** Western blot quantification of agonist-induced AKT S473 phosphorylation. Data was normalized to phospho-AKT S473:total AKT and to 5 min high plasma membrane expression sample. **e** Representative western blot illustrating CXCL12-induced ERK1/2 phosphorylation in high and low plasma membrane expressing CXCR4 cells. **f** Western blot quantification of CXCL12-induced ERK1/2 phosphorylation. Data was normalized to phospho-ERK/12:total ERK1/2 and to 5 min high plasma membrane expression sample. All experiments were conducted in RPE cells overexpressing WT CXCR4 and stimulated with 12.5 nM CXCL12 for the stated time course. Individual data points from each experiment are plotted; mean, SD, median line. Statistical significance (*) denotes $p < 0.05$.
